# Targeting lignocellulolytic gene clusters in novel *Trichoderma atroviride* and *Trichoderma harzianum* strains through Bac-guided analysis

**DOI:** 10.1101/2023.09.28.559926

**Authors:** Paulo Henrique Campiteli, Maria Augusta Horta, Rafaela Rossi Rosolen, Juliano Sales Mendes, Carla Cristina da Silva, Danilo Sforça, Anete Pereira de Souza

## Abstract

Lignocellulosic biomass is known as a challenging substrate for enzymatic hydrolysis, increasing the processing cost in biorefineries. In nature, filamentous fungi, including those of the genus *Trichoderma*, naturally degrade lignocellulose by using an arsenal of hydrolytic and oxidative enzymes that act synergistically with biomass degradation. This work explored the genome organization of target genes of *Trichoderma atroviride* and *Trichoderma harzianum* that are able to promote hydrolysis by identifying regions enriched in degradative enzyme-encoding genes, namely, hydrolytic clusters. We employed bacterial artificial chromosome (BAC) methodology to target specific genomic regions to explore their genetic organization, proximal gene context, and gene expression under degradative conditions. Using this tool, it was possible to inspect the linear structure and expression data of target hydrolytic-rich genomic regions. The results offered a genomic perspective of the organization of genome regions related to carbohydrate metabolism, revealing novel genome regions and genes that are positively regulated during cellulose degradation and contributing to elucidating differences in gene organization among *Trichoderma* species.

**Highlights:**

- High-quality genomic regions were selected using closely related species CAZyme targets.
- RNA-seq data were used to quantify BAC genes expression.
- Relevant regions were identified by combining functional and differential expression data.
- Novel targets for future characterization identified via differential expression.

## Introduction

*Trichoderma* spp. (ascomycetes) are opportunistic generalists that are known to be efficient plant biomass decomposers (Druzhinina et al., 2018; Kubicek et al., 2019). They are distributed worldwide and colonize soil, living plants, decomposing wood, and other fungi (Druzhinina et al., 2018; Kubicek et al., 2019; Srivastava and Shahid, 2014; Zeilinger et al., 2016). These species have previously been of interest in biotechnology due to their efficient lignocellulolytic activities, especially *Trichoderma reesei* QM6a (and its derivative clones) (Li and Wang, 2021; Li et al., 2017). They are widely used for the industrial production of cellulases and hemicellulases for enzymatic cocktails, which are applied in saccharification (Bischof et al., 2016) and are used to optimize enzymatic hydrolysis processes in biorefining, especially during the process of lignocellulose biodegradation. Biodegradation promoted by *Trichoderma* spp. is driven by a complex network of reactions, which first demands substrate perception, triggering specific metabolic reactions to synthesize and transport hydrolytic enzymes, classified as carbon-active enzymes (CAZymes), and other genes related to lignocellulolytic activity, such as transcription factors (TFs), transporters, and other complex metabolites (Andlar et al., 2018). CAZymes are enzymes related to cleavage, synthesis, or interactions with carbon polymers and are organized and classified into families and subfamilies according to the reactions in which they participate as shown in the Carbon Active Enzymes database (http://www.cazy.org/) (Andlar et al., 2018; Drula et al., 2022).

In addition to its potential as a source of new and optimized hydrolytic enzymes, emerging interest in *Trichoderma* is linked to the remarkable biological activities of these species, with their outstanding biocontrol and biofertilization capacities. Some species are capable of controlling rhizospheric microorganisms while displaying beneficial interactions with plants and are considered prominent biocontrol agents (BCA) that pose low risk to humans (Kubicek et al., 2019; Sood et al., 2020). *Trichoderma atroviride* and *Trichoderma harzianum* are mycoparasites that exhibit biocontrol activity against various phytopathogens, including *Rhizoctonia* sp., *Botrytis* sp., *Fusarium* sp., and *Penicillium* sp. (Nascimento et al., 2022; Sood et al., 2020; Stracquadanio et al., 2020). Additionally, plant-*Trichoderma* cocultures exhibit biofertilization activities, promoting vegetative traits and immunologic defense systems and attenuating abiotic stress (Guzmán-Guzmán et al., 2023; Kubicek et al., 2011, 2019; Sood et al., 2020; Tyśkiewicz et al., 2022).

By observing the capabilities of the species of the genus *Trichoderma*, it is possible to conclude that Trichoderma possesses a remarkable system of substrate perception and that its members are able to trigger specific reactions for metabolic adaptation by synthesizing and transporting proteins and complex metabolites for substrate degradation, defense, competition, and interaction with other organisms (Andlar et al., 2018). To explore the diversity in the genus and the differences in genome organization and CAZyme composition, we investigated *T. harzianum* CBMAI 179 (TH0179) and *T. atroviride* CBMAI 0020 (TA0020). Both species were previously investigated under biomass degradation conditions. The hydrolytic potential of these strains was evaluated by quantifying cellulase and hemicellulase enzymatic activities (Horta et al., 2018), and the transcriptomic profiles of both strains were explored by gene expression analysis under cellulose degradation conditions (Almeida et al., 2021). TH0179 exhibited increased enzymatic activity under cellulose degradation conditions, while TA0020 exhibited a more complex transcriptome profile with decreased lignocellulolytic activity (Almeida et al., 2021; Horta et al., 2018; Rosolen et al., 2022). The observed heterogeneous responses to cellulose degradation indicate the high complexity of gene expression at the genus level, in which closely related species can carry relevant species-specific traits that affect the biological response of the fungus.

Bacterial artificial chromosome (BAC) methodology is a molecular tool used to recover and study target genomic regions (Peterson et al., 2000). It is a selection platform that distributes the entire genome in libraries that allows the identification of specific genomic region targets by PCR. This technique can be applied for gap closure in fragmented assemblies, as an assembly method (as described for the first human genome sequence (Waterston et al., 2002)), or as a tool for studying targeted genomic regions in higher eukaryotes with complex genomes rich in polyploid and aneuploid repetitive elements (Mancini et al., 2018; Visendi et al., 2016). It is a simple, reliable, low-cost method for studying target genomic regions and is especially powerful for studying novel species without previous genomic data. In fungi, the BAC approach has been successfully applied to investigate genomic regions with CAZyme cluster genes in *T. harzianum* IOC 3844 (Crucello et al., 2015; Filho et al., 2020). In *Streptomyces* spp., for example, the BAC approach was used for genome mining and revealed important gene clusters of bioactive natural products and the corresponding biosynthetic pathways (Xu et al., 2016). Another study used BAC libraries to characterize a novel gene conferring sensitivity to the necrotrophic fungal pathogen *Parastagonospora nodorum* (Zhang et al., 2021).

In this study, the BAC approach was applied to investigate hydrolytic-related genomic regions in the TH0179 and TA0020 strains. This methodology allowed us to describe genomic regions enriched in coding cellulases, hemicellulases, and ligninases, as well as TFs, transporters, and potential targets involved in cellulose degradation. In addition, we employed the data generated from a previous RNA-Seq study (Almeida et al., 2021) to analyze the expression of CAZyme genes through a BAC-guided analysis, which allowed the identification of lignocellulolytic CAZyme clusters that are positively regulated for the degradation of cellulose.

The results offer insights into the genomic organization and gene clustering of hydrolytic enzymes in the studied strains and were integrated with the transcriptomic data. The results addressed the differences in lignocellulolytic activities between species. Additionally, species-specific genomic regions and gene targets were described for future characterization.

## Materials and methods

### Strains and reference sequences

TH0179 and TA0020 were obtained from the Brazilian Collection of Environment and Industry Microorganisms (CBMAI), located in the Multidisciplinary Center for Chemical, Biological, and Agricultural Research (CPQBA), Campinas, São Paulo, Brazil. The identities of the *Trichoderma* isolates used in these studies were authenticated by a provider based on phylogenetic studies of their internal transcribed spacer (ITS) region, translational elongation factor 1 (tef1) marker gene and RNA polymerase II (rpb2) marker gene (Kimura et al., 1980; Raeder et al., 1985; Saitou et al., 1987; Tamura et al., 2007; Thompson et al., 1994). The phylogenetic tree based on the ITS region of this species was previously published (Rosolen et al., 2022).

As the genomes of the studied species were not available at the time of the experiments, *T. harzianum* T6776 (Baroncelli et al., 2015) was used as a reference genome for TH0179, and *T. atroviride* IMI206040 (Kubicek et al., 2011) was used as a reference genome for TA0020. The reference genomes provided the consensus sequences for the target gene primer design and for RNA-Seq read mapping. The transcriptomes of TH0179 and TA0020, in which fungi were cultivated with crystalline cellulose (inducing degradative conditions) or glucose (control conditions) as a unique carbon source, were previously described (NCBI <u>BioProject</u> <u>PRJNA336221</u>) (Almeida et al., 2021).

### BAC library construction and screening

For each species, 10 µl of spore culture was inoculated on potato dextrose agar (PDA) plates (100 µg.ml-1 ampicillin, 34 µg.ml-1 chloramphenicol) at 28°C for four days. Next, a 2 cm × 2 cm portion of the colony was inoculated on sterilized potato dextrose and incubated (28°C, 3 days, 200 rpm). The final liquid culture was filtered using sterile *Miracloth* (Calbiochem, USA), and the mycelia were subjected to high-molecular-weight genomic DNA (HMW-gDNA) extraction using the Qiagen DNeasy Plant Mini Kit II (Hilden, Germany). The HMW-gDNA was immobilized on low-melting agarose plugs for BAC library construction, and large genomic DNA inserts were cloned and inserted into *E. coli* using the pAGI BAC vector (provided by The French Plant Genomic Resources Center, CNRGV, INRAE, France). Inserts were transformed using *E. coli* DH10B competent cells by electroporation.

The transformed culture was inoculated on PDA plates, and the successfully inserted fragments were identified visually as white colonies. Each individual colony represented a unique clone and was cultured and plated on 384-well plates. Each library was examined on a rapid selection platform composed of combined rows or column pools, which allowed fast screening of the library through real-time qPCR of target genes using a previously described method (Crucello et al., 2015; Filho et al., 2020). Supplementary material S1 provides the primer sequences designed based on target CAZymes (Untergasser el al., 2012) and the clone localization in the library. The entire BAC library consisted of 5,760 individual clones, each carrying a unique long genomic fragment (20 kb to 200 kb), constructed following (Peterson et al., 2000), with minor modifications for filamentous fungal samples.

### Screening of CAZyme genes and BAC-DNA extraction

The primers were used for amplification on the selection platform, and BACs were selected via melting curve comparison. To avoid nonspecific and false positive amplification, the final selection was performed by 3% agarose electrophoresis (80 V, 2 h, 1 kB Promega ladder). The selected clones were preincubated in growth medium (15 ml, 200 rpm, 37°C, 6 h-12 h) and then incubated overnight in 80 ml of growth medium (200 rpm, 37°C, 6 h-12 h). After incubation, the culture was centrifuged, and BAC-DNA was extracted using a Marcherey-Nagel Plant Mini Kit II (Düren, Germany) following standard procedures.

### BAC-DNA assembly and evidence-based gene prediction and expression

The selected BACs were sequenced by the Arizona Genomics Institute (Pacbio Sequel II, Arizona, USA). The sequencing data were transferred to a local server for polishing and assembly. The raw reads were polished using Quiver, and the BACs were *de novo* assembled using Canu v.2.1.1 (Koren et al., 2017). The assembled genomic regions were aligned to the *T. harzianum* T6776 (Baroncelli et al., 2015) and *T. atroviride* IMI 206040 (Kubicek et al., 2011) reference genomes to identify *Trichoderma* target sequences from the respective bulk BAC sequences (**Supplementary Material 1**). BACs <40 kb were filtered out of the analysis. Then, all-vs.-all alignment was performed for the TA0020 and TH0179 BACs individually to select nonoverlapping regions. For the structural comparison, BACs were aligned to the *T. atroviride* IMI 206040 (Kubicek et al., 2011) and *T. harzianum* CBS226.95 (Baroncelli et al., 2015) draft genomes and the *T. reesei* QM6a (Li et al., 2017) reference genome using MUMmer (Marçais et al., 2018). The MUMmer dot plot parameters employed to visualize the alignment were as follows: all-vs.-all, minimum match 50 bp, and minimum alignment length 100 bp.

For evidence-based gene prediction, we first aligned RNA-Seq reads to the selected BACs using STAR (Dobin et al., 2013). BAM outputs were input into StringTie 2.2.1 (Pertea et al., 2015) for transcript assembly guided by the BAC sequences. The final set of transcripts was used as input for maker2 2.31.11 (Holt and Yandell, 2011) gene structure prediction. The predicted models were used for further functional inference and comparative analysis. Gene models were processed with InterproScan v5.55-88.0 (Jones et al., 2014; Quevillon et al., 2005) and eggNOG-mapper v.2.0.1 (Huerta-Cepas et al., 2019). Parallel annotation of modules and clusters coding CAZymes was conducted with the dbCAN v3 web platform (Zheng et al., 2023). KOALAblast (Kanehisa et al., 2016) was used to identify metabolism-related genes from http://www.kegg.jp (Kanehisa and Goto, 2000) via a search against *Hypocreceae*. The antiSMASH web server 6.0 (Blin et al., 2021) was used to define secondary metabolite biosynthesis-related gene clusters (SMBGCs). Functional data and gene structure predictions were integrated using Funannotate (Li and Wang, 2021; Palmer and Stajich, 2020) implemented on the Galaxy platform (Galaxy Community, 2022).

The expression values of genes present in the genomic regions of TH0179 and TA0020 selected by BAC-guided assembly were extracted from previously published data (Almeida et al., 2021) using FastQC v0.11.5 (Trivedi et al., 2014) and Trimmomatic 0.39 (Bolger et al., 2014) parameters as follows: a minimum length of 36 bp and a SLIDINGWINDOW ratio of 4:10:10 were used to access and filter the reads by quality. Filtered reads were aligned to BAC sequences on a STAR 2.7.10b aligner (Dobin et al., 2013), and StringTie 2.2.1 (Pertea et al., 2015) was used for transcript assembly. In addition, the TPM value for each gene was calculated automatically based on StringTie quantification. The EdgeR package (Robinson et al., 2010) was used to compare the gene expression under the studied conditions and identify genes with significant differential expression.

To define orthologous genes, OrthoFinder v2.5.2 (Emms and Kelly, 2019) compared newly annotated gene models among 24 *Trichoderma* species, with *Saccharomyces cerevisiae* S288C (Goffeau et al., 1996), *Neurospora crassa* OR74A (Galagan et al., 2003), *Pleurotus ostreatus* PC15 (Riley et al., 2014), and *Phanerochaete chrysosporium* RP-78 (Ohm et al., 2014) as external groups, all of which were extracted from the mycocosm (Grigoriev et al., 2014) (the list of species is available in **Supplementary material 2**). The comparative genetics and per-species outputs of OrthoFinder are available in **Supplementary material S2**.

## Results

### Target BAC selection

The final BAC library consisted of 5,760 individual clones, each carrying a unique long genomic fragment (ranging from 20 kb to 200 kb), and was examined using a rapid selection platform with plates containing pools for rapid screening of the whole library. For genomic region screening, (I) 42 and (II) 54 BAC clones were selected for TH0179 and TA0020, respectively (primers available in Supplementary material 1). Each positively extracted clone was subjected to quality control, and the profile of the pooled samples integrity was checked via a fingerprint test (HindIII BAC digestion) involving 1% agarose electrophoresis and 3% agarose electrophoresis. After BAC sequencing, all-vs.-all sequence alignment was performed for each strain. Only BACs larger than 40 kb were considered for further analysis. After filtering, a final set of 24 nonredundant genomic regions for TH0179 and 44 for TA0020, ranging in size from 41 kb to 138 kb and 53 kb to 164 kb, respectively, was obtained. BACs were submitted to NCBI via BankIt as Bioproject PRJNA1028979 Bio samples TH0179 SAMN37846728 and TA0020 SAMN37846658. The table 1 presents an overview of the achieved numbers during BAC characterization and functional annotation.

**Table 1.**
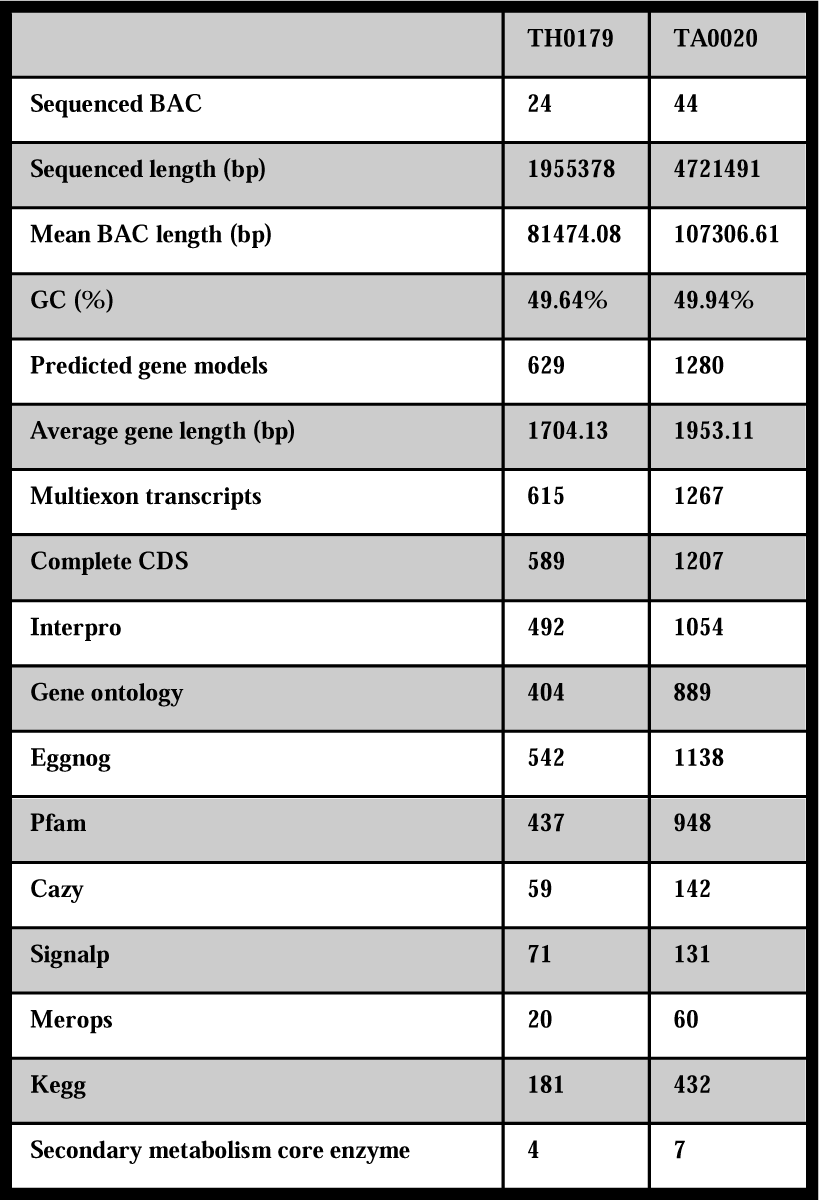
Overview of the BAC genomic characterization and functional annotation of the evaluated strains.

The TH0179 genomic region represents 1.9 Mb, ∼5% of the *T. harzianum* CBS226.95 genome. On the other hand, 44 unique regions were recovered from TA0020, reaching 4.7 Mb, 15% of the *T. atroviride* IMI 206040 genome. A comparative analysis of the generated sequences with reference genomes showed that the alignment rate compared to T*. reesei* QM6a was 12.3% for TH0179 and 7.5% for TA0020. A total of 95.5% of TH0179 was aligned to the *T. harzianum* CBS226.95 reference, and 87.3% of TA0020 was aligned to TaIMI 206040 as a reference. Notably, there is variation in the gene position among species, and Figure 1 shows the distribution of the aligned sequences in the different reference genomes, with the presence of orthologs in different genomic regions, despite a tendency toward similar gene organization that increases with species similarity.

**Fig. 1.**
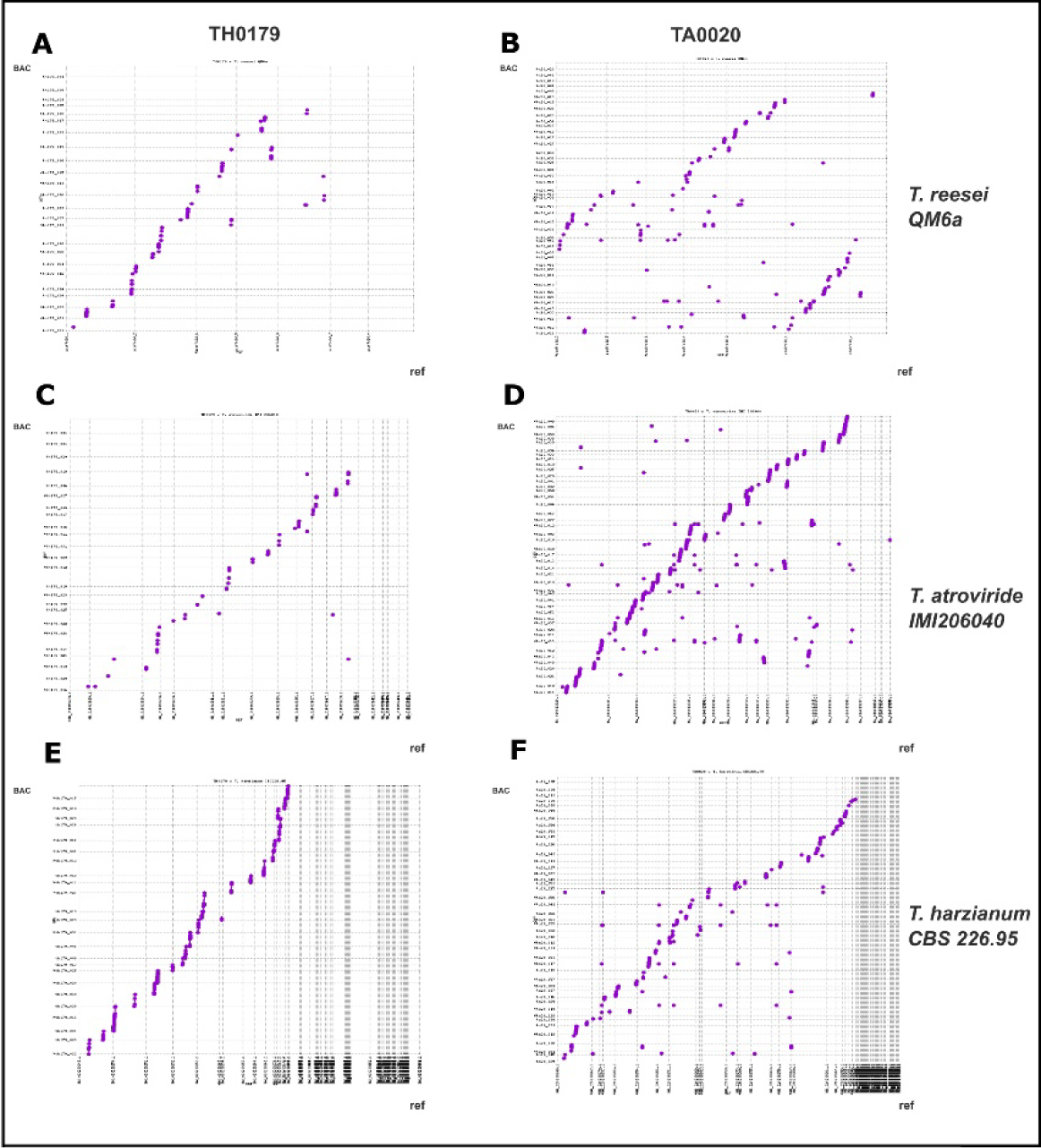
MuMmer alignment to *Trichoderma* references. Alignments visualized using a MUMmer dot plot with the parameters Minmatch 50 bp and match length 500 bp against *T. reesei* QM6a, *T. harzianum* CBS22695 and *T. atroviride* IMI 206040. (a) TH0179 x *T. reesei* QM6a. (b) TA0020 x *T. reesei* QM6a. (c) TH0179 x *T. atroviride* IMI206040. (d) TA0020 x *T. atroviride* IMI206040. (e) TH0179 x *T. harzianum* CBS226.95 (f) TA0020 x *T. harzianum* CBS226.95.

To establish orthologous relationships, we analyzed 30 fungal species plus the gene models generated in this study. In total, 310,808 genes were analyzed, 94.6% of which were assigned to 19,020 orthogroups, including 1,392 species-specific orthogroups. Forty-eight orthogroups were shared among all of the strains (Supplementary Material 2). A total of 1,343 (92.2%) TA0020 genes were assigned to orthogroups, with 37 genes remaining unassigned, and they were distributed in almost all 24 genomic regions. Among the TH0179 genes, 596 (94.8%) were assigned to orthogroups, and 33 remained unassigned and distributed in 17 genomic regions. In addition, 168 orthogroups were shared among TH0179 and TA0020, including genes encoding 15 CAZymes, 12 TRs, 10 TFs and 5 SMBs.

Multiple tools were used to access functional data for the annotated models. Ninety-three percent of the TA0020-annotated genes and 91% of the TH0179-annotated genes had at least one functional annotation. CAZyme modules represented 9.3% of the TH0179 annotated gene model and 11% of the TA0020 annotated gene model (Table 1, Fig. 2, Supplementary material 3).

**Fig. 2.**
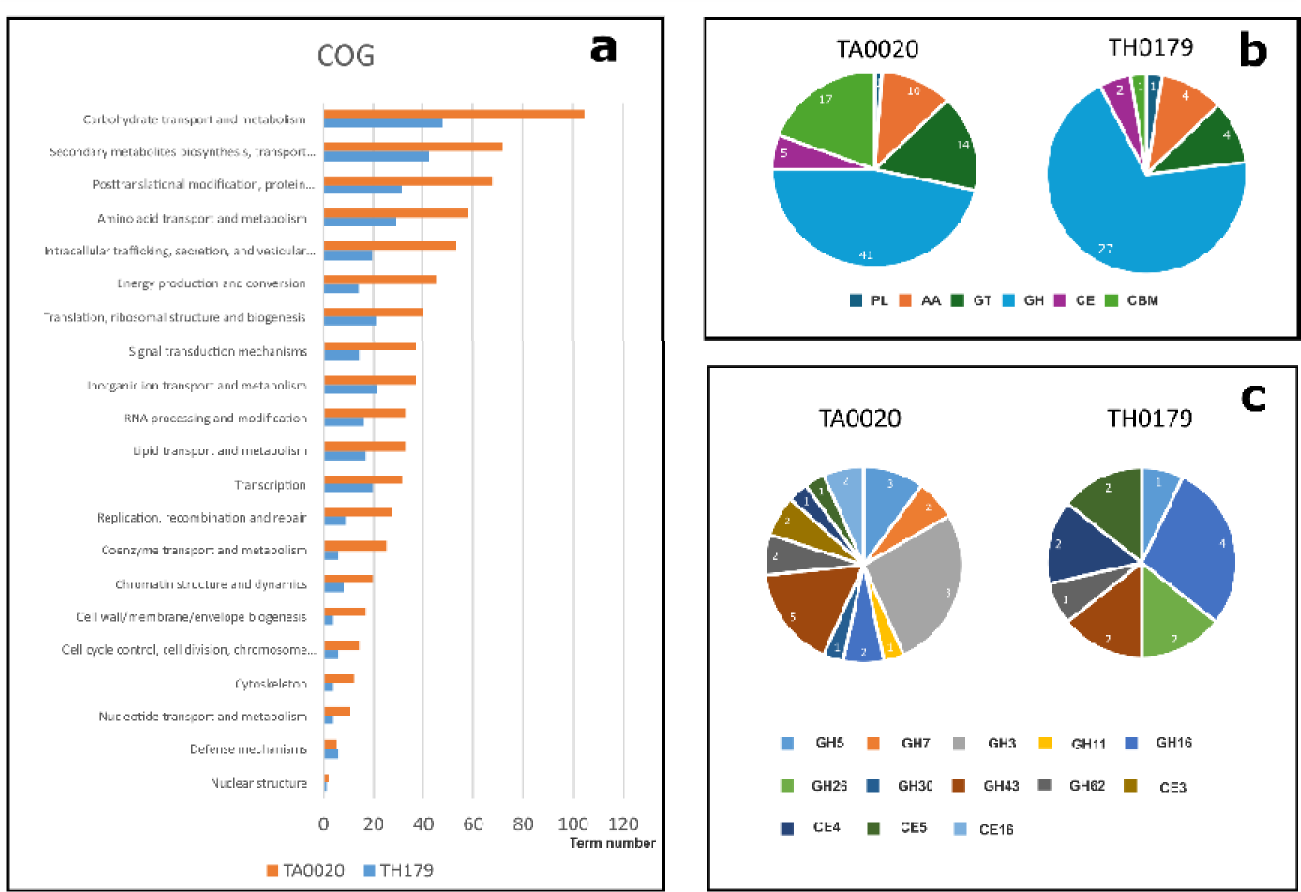
BAC functional inference. (a) COG categories attributed to annotated BAC genes. (b) CAZyme families annotated in each strain. (c) Lignocellulolytic CAZymes (Andlar et. al 2018) annotated for each strain. The numbers of classified genes for each class are shown in the plot.

### CAZymes, transporters and transcription factors and clusters

In total, 59 CAZymes, 27 TFs, and 39 TRs were recovered from TH0179. Seventy-one genes presented signal peptides, including 23 CAZymes, one TR, one TF, and two SMGs, that were linked to the NRPS core enzyme from OR633216. Forty-two percent of the annotated CAZymes had predicted substrates, four of which had multiple targets. Regarding CAZyme clustering, in TH0179, three genomic regions did not present any CAZymes, and 5 regions presented only one each. The other 15 selected regions had 2 to 6 CAZymes, among which OR633232 was considered the BAC with the AA2, CE4, GH5, CE4, and AA11 CAZymes; interestingly, all of these were to some extent related to cellulose degradation, with the GH and CE genes considered lignocellulolytic, while the AA redox family is involved in complex substrate degradation. DBcan3 can also be used to identify signature CAZyme clusters based on genomic localization and homology to known clusters in the database. For TH0179, 14 regions showed annotated CAZyme clusters (Supplementary material 3).

For TA0020, 143 CAZymes, 62 TFs, and 114 TRs were identified. A total of 131 genes presented signal peptides, including 43 CAZyme, two TF, two TR and five SM genes. Forty percent of the CAZymes of TA0020 had predicted substrates, with 13 showing more than one target substrate. Regarding CAZyme clustering, four regions did not present any CAZymes, four regions presented one CAZyme each, and 36 genomic regions presented two or more CAZymes. Additionally, 31 regions contained dbcan3-annotated CAZyme clusters (Supplementary material 3).

Overall, the TA0020 gene models presented a greater number of annotated CAZymes, TFs and TRs than did TH179, but for both strains, it was possible to identify regions in which those hydrolytic genes are near the same genome position, in tandem with other genes predicted to be involved in metabolic hydrolysis control.

### Expression analysis under cellulose degradation conditions

For all BACs, 62 genes were significantly differentially expressed (they were DEGs), 35 of which were upregulated when cellulose was used as a carbon source for culture and 27 of which were downregulated (**Supplementary material 3**).

The downregulated DEGs included GH17 (TA0020_0933), AA9 (TA0020_1125), CBM18 (TA0020_1121), three TRs (TA0020_0935, TA0020_0744, TA0020_1208), 6 TFs (TA0020_0369, TA0020_0374, TA0020_0371, TA0020_0358, TA0020_0735, TA0020_1197), an additional biosynthetic SM gene from the T1PKS SMBGC cluster (TA0020_0056), and a direct neighbor gene from the Terpene core-enzyme SMBGC (TA0020_1004). The cellulose DEGs included GH34 (TA0020_0446), GT1 (TA0020_1099), GT20 (TA0020_1261), GH28 (TA0020_0170), three TFs (TA0020_1100, TA0020_0593, TA0020_0023), two genes from the Ripp-like SMBGC, one with no annotation (TA0020_0494) and one with an alpha-beta hydrolase fold domain (TA0020_0489). Among the DEGs, the functional inference for two downregulated and five upregulated genes remained unclear, and these genes are therefore potential new targets for future characterization.

Among the 14 DEGs identified in all of the TH0179 BACs, three were downregulated, and 11 were upregulated; these DEGs were distributed in 9 genomic regions. The downregulated genes were TH0179_0577 (eggNOG hit with a hypothetical protein ortholog), TH0179_0308 TF (Zn(2)-C6 fungal type) and TH0179_0309 TR (MSF superfamily). The 10 upregulated genes included three CAZymes: TH0179_0102 (GH18), TH0179_0045 (AA7) and TH0179_0001 (GH95). Additionally, TH0179_0393 and additional biosynthetic genes from SMBGC, TH0179_0359 (tetratricopeptide and protein protein interaction driver), TH0179_0589 (WSC domain, a carbohydrate binding domain), TH0179_0302 (PAN domain), TH0179_0217 (only an eggNOG hypothetical protein hit), TH0179_0053 TR (related to cation transport), and TH0179_0248 (CorA-like Mg2+ transporter protein), were identified. AA7 also belongs to an annotated CAZyme cluster in the OR633223 genomic region represented in the next section (Fig. 3).

**Fig. 3.**
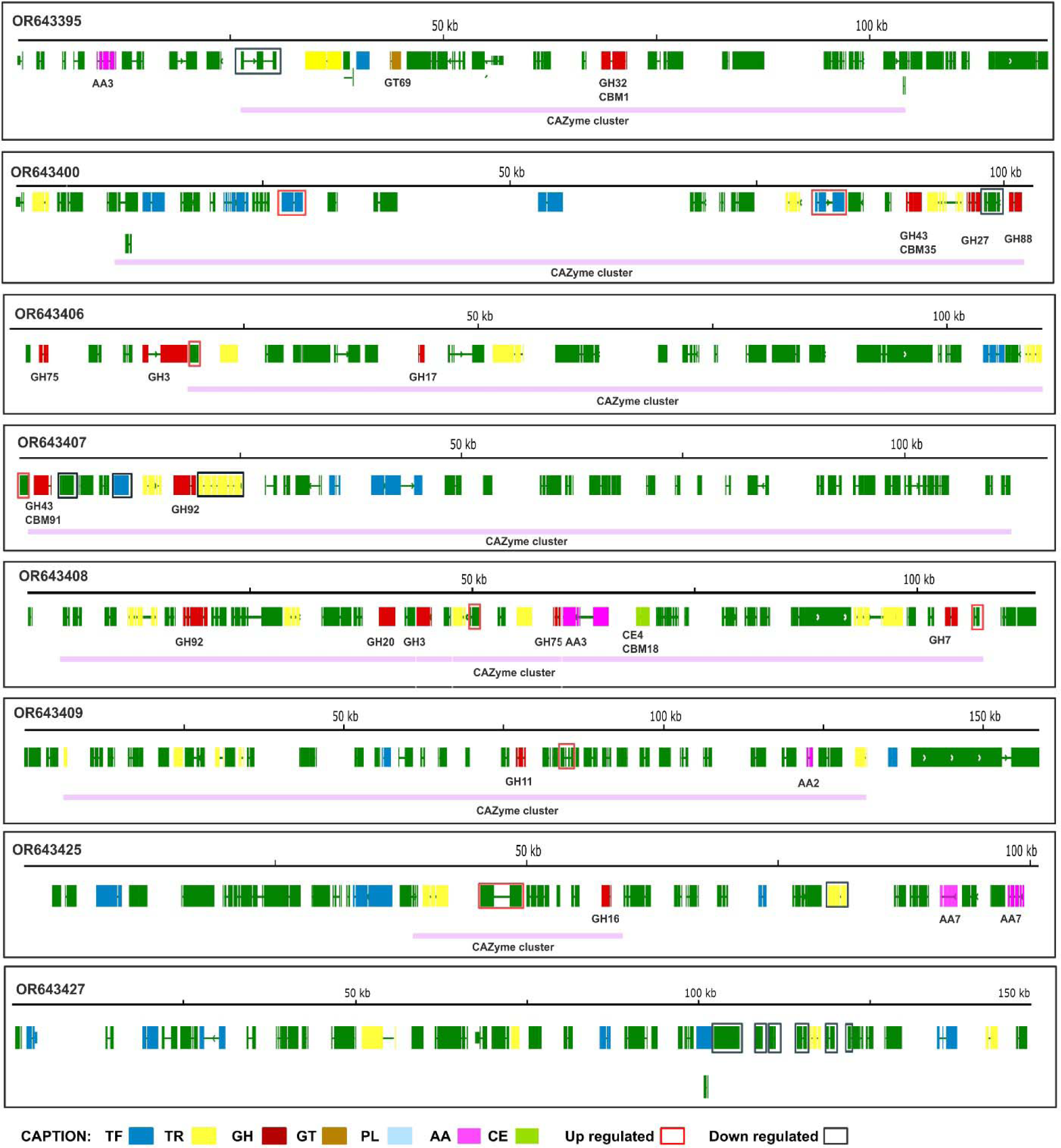
TA0020 CAZyme clusters or regions related to cellulose and glucose degradation. TA0020 IGV visualization of CAZyme clusters or DEG-carrying regions identified via functional inference, genomic context, and differential expression analysis.

### Major degradative regions and clusters

For in-depth investigation of the genome regions captured by the BACs, the gene clusters that presented degradative genes were modeled, and the gene position, annotation and/or expression information is shown in Fig. 3 for TA0020 and Fig. 4 for TH0179.

**Fig. 4.**
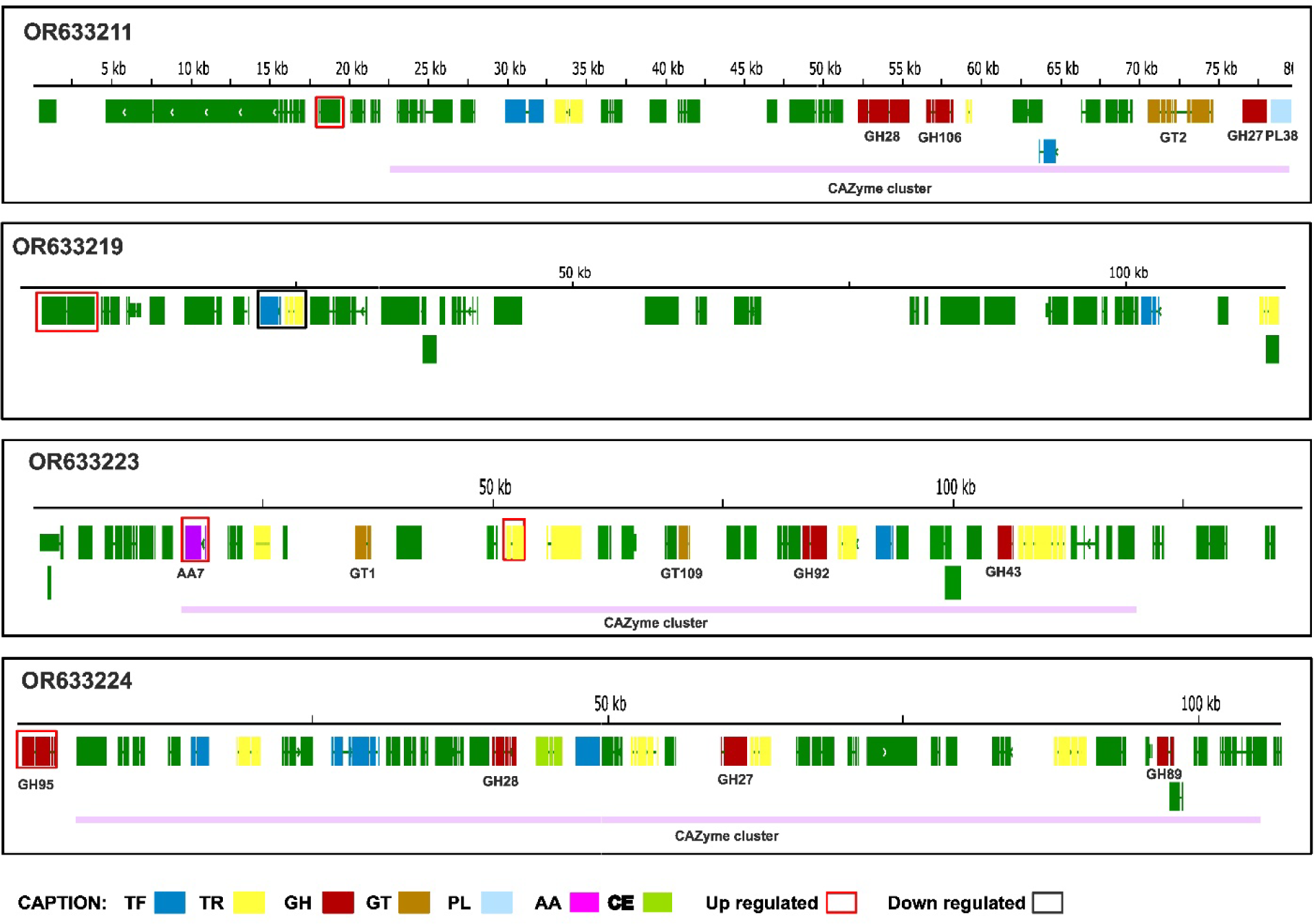
TH0179 CAZyme clusters or regions related to cellulose and glucose degradation. Major TH0179 IGV visualization of CAZyme clusters or DEG-carrying regions identified via functional inference, genomic context, and differential expression analysis.

By crossing the DEGs and the CAZyme clusters, it was possible to identify major cellulolytic and glycolytic CAZyme clusters among the annotated BACs. For TA0020, eight relevant genomic regions were identified

OR643395 carries three CAZymes. The first gene inside the annotated CAZyme clusters was downregulated TA0020_0615 (a flavin reductase related to redox activities). OR643400 carries three CAZymes and multiple TFs and TRs. Within the cluster, two TFs, TA0020_0737 and TA0020_0745, which are both Zn(2)-C6 fungal-type transcription factors, were upregulated. One gene was downregulated TA0020_0751 (alginate lyase). OR643406 also carries three CAZymes; the first gene from the CAZyme cluster, TA0020_0927 (pyridoxal phosphate-dependent enzyme), a direct neighbor of GH3, was upregulated. OR643407 carries two CAZymes, both inside the annotated CAZyme cluster. TA0020_0445 (ferric reductase NAD binding domain) and TA0020_0473 (no functional hit) were upregulated. In contrast TA0020_0447 (carboxylesterase family, BD-FAE a novel bifunctional CE), TA0020_0450 TF (fungal Zn(2)-Cys(6)) and TA0020_0453 TR (ABC transporter and transmembrane region domain) was downregulated. OR643408 carries 6 CAZymes, all of which are inside the annotated CAZyme cluster. Four genes were upregulated: TA0020_0234 (GH20), TA0020_0239 (taurine catabolism dioxygenase TauD, TfdA family), TA0020_0251 TR (peptidase merops family M28, MSF superfamily) and TA0020_0252 (taurine catabolism dioxygenase TauD, TfdA family). OR643409 carries two CAZymes, both inside the CAZyme cluster. Two genes were upregulated: TA0020_0203 (amino-transferase class IV) and TA0020_0211 (amidohydrolase family). OR643425 carries a CAZyme (GH26) inside the annotated clusters, plus two others in the region’s border. Inside the clusters, the gene TA0020_0678 (fungal pheromone mating factor STE2 GPCR and Indoleamine 2,3-dioxygenase domains) was upregulated. Outside the cluster, the gene TA0020_0689 TR (MSF superfamily) was downregulated. OR643427 did not present any CAZyme. Also, seven genes were down regulated. One was TA0020_0380 (amino acid permease). Interestingly, the other six downregulated genes belonged to the annotated SMBGC, including the NRPS-like core biosynthetic enzyme TA0020_0395 (AMP-binding enzyme, phosphopantetheine attachment site, transferase). TA0020_0396 (pyoverdine/dityrosine biosynthesis protein, sporulation related), TA0020_0397 (enoyl-(acyl carrier protein) reductase), TA0020_0398 (male sterility protein), TA0020_0400 (2OG-Fe(II) oxygenase superfamily, nonheme dioxygenase in morphine synthesis N-terminal), and TA0020_0401 (DUF218 domain).

For TH0179, 9 regions presented DEGs, among which 3 contained CAZyme clusters with upregulated genes. One region without CAZymes but carrying three DEGs was identified.

OR633211 carries five CAZymes, all inside the annotated CAZyme cluster. In addition, one SMBGC was identified in the same region. The expression of the gene TH0179_0393 (zinc-binding dehydrogenase), which belongs to the SMBGC, was upregulated. OR633219 carries no CAZyme. The expression of the TH0179_0302 gene (PAN domain, which mediates binding to high-molecular-weight (HMW) kininogen) was upregulated. TH0179_0308 TF (Zn(2)-C6 fungal-type) and TH0179_0309 TR (MSF superfamily) were downregulated. OR633223 carries a CAZyme cluster with 5 CAZymes, and interestingly a SMBGC overlaps in the region. TH0179_0045 (AA7), which belongs to both functional clusters, and TH0179_0053 TR (haloacid dehalogenase-like hydrolase) were upregulated. OR633224 carries 4 CAZymes, three inside the annotated cluster. TH0179_0001 (GH95), which is not part of the cluster, was upregulated.

Overall, relevant genomic regions with CAZyme-annotated genes and clusters were observed. The expression analysis of TH0179 and Ta0020 helped to identify major players in degradative conditions that are likely active to respond to the current conditions, and which are also integrated with other genes in their vicinities.

## Discussion

BAC-guided transcriptome analysis was used to mine high-quality genomic regions from TA0020 and TH0179 containing CAZyme targets with biotechnological relevance. This methodology promoted *hybrid* evidence-based and *ab initio* gene prediction, followed by functional inference and alignment-based expression analysis to identify regions potentially linked to cellulose and glucose degradation. Overall, the main goal was to inspect the genomic and transcriptomic variations and review the genomic context of potential CAZymes and other relevant features related to the cellulolytic metabolism process. A range of clustered CAZymes, including cellulases, hemicellulases, and ligninases, were identified. Additionally, important genes, such as SMBGC, oxidases, proteinases, TFs and TRs, were closely related to CAZymes in the gene sequence clusters. Additionally, genes with high expression levels under degradative conditions were identified and are likely relevant to the degradative process. The exact relationships of the clustered genes, how their expression is orchestrated to respond to lignocellulose synergistically, and the species-specific particularities affecting the ligninolytic process were reviewed from a nonmodel species perspective of *Trichoderma* species.

### Genomic region characterization

The BAC approach is a methodology that can successfully target specific regions and guarantee the quality and stability of specific genomic regions; thus, this method avoids assembly mistakes, especially in repetitive regions of the genome. This work explored specific regions of the genome; however, the total length obtained for TA0020 was greater than that for TH179, and the differences in the number of genes identified between the two strains can be explained by this limitation (Fig. 2). Therefore, the BAC sequences obtained and compared with the genomes reinforce the diversity among the genus (Fig. 1). Previous comparative analysis revealed that 50-25% of the *Trichoderma* genes are species-specific and not shared with other *Trichoderma* species. Additionally, gene families such as ankyrin, heterokaryon incompatibility domain protein (HET), TF, carbohydrate degradative, and SMBGC genes all present high polymorphism rates (Kubicek et al., 2011, 2019; Mukherjee et al., 2013). The intraspecific diversity of the *Harzianum* clade is a challenge to modern molecular genetics because this species aggregates multiple highly complex infra groups (Druzhinina et al., 2010; Kubicek et al., 2019; Schalamun and Schmoll, 2022), which can partially explain the limited CAZyme gene transfer.

### Comparative analysis

The BAC alignment with the reference genome and orthology data helped in the assessment of the complexity of *Trichoderma* at the strain level. *T. reesei* QM6a is considered a model *Trichoderma* species that is phylogenetically distant from other biocontrol species (Kubicek et al., 2019). As expected, the BAC correspondence decreased with phylogenetic distance. Based on the alignment to its own species reference (Fig. 1), it is possible to conclude that TA0020 may be more distant from *T. atroviride* IMI206040 than TH0179 is from *T. harzianum* CBS226.95. Additionally, for all three comparisons, the TA0020 alignments showed greater complexity, with the same BAC aligning to multiple contigs/chromosomes in the references. In the orthology analysis, only 33 TH0179 genes were unassigned to orthogroups, while 37 genes for TA0020 remained unassigned. According to a previous molecular phylogenetic analysis of rpb2, *T. atroviride* species belong to a basal branch that diverged early from other species, most likely maintaining characteristics of a common ancestor (Kubicek et al., 2019).

Overall, the analysis of the BAC gene regions corroborated the findings of rapid evolution rates due to the opportunistic lifestyle of *Trichoderma*. The rapid evolution of the genus reshaped their genomes, allowing them to colonize potential niches (Kubicek et al., 2019). Their remarkable intraspecific capacities are greatly influenced by lateral gene transfer and other complex evolutionary processes contributing to genomic variations, even those observed in closely related species. Interestingly, carbohydrate metabolism and biocontrol gene families seem to be especially influenced by evolutionary processes (Guzmán-Guzmán et al., 2023; Hewedy et al., 2020; Kubicek et al., 2011, 2019; Schalamun and Schmoll, 2022; Sood et al., 2020; Tyśkiewicz et al., 2022).

### CAZyme genes and clusters

CAZymes play a central role in carbon substrate degradation, energy consumption, cell wall synthesis, and other accessory activities mediated by substrate availability and other environmental stimuli (Drula et al., 2022).

For the studied species, the retrieved CAZyme genes represent a tiny fraction of all CAZymes (Filho et al., 2017; Kubicek et al., 2011; Rosolen et al., 2023). Whole-genome analysis has shown that *T. reesei* genomes contain fewer CAZymes than do those of biocontrol strains (Kubicek et al., 2011; Li and Wang, 2021; Li et al., 2017). This is related to the fact that *T. reesei* is a saprophytic species that has lost biocontrol-related gene families, including important CAZyme genes (Kubicek et al., 2019), and the recent whole-genome comparative analysis of *T. harzianum* IOC 3844, TH0179, and TA0020 and *T. reesei* 711 corroborated these differences (Rosolen et al., 2023). For the retrieved genomic regions in the present study, TA0020 showed a diverse set of cellulases, hemicellulases and ligninases despite its lower lignocellulolytic potential and lower expression of CAZyme genes, which may be related to plant-*Trichoderma* interactions partially degrading the plant cell wall, allowing the exchange of nutrients in endophytic species.

Complex substrate degradation, such as that of glucose, cellulose, lignocellulose, and chitin, activates different and complex synergistic reactions with different sets of genes, including CAZymes, TFs, and TRs. The clustered distribution of genes contributing to the same phenotype is not new, but in eukaryotes, genes are expressed individually, and the clustering of metabolic genes in the fungal genome is still unclear (Rokas et al., 2018; Slot, 2017; Wisecaver and Rokas, 2015). Regarding the clustered distribution of CAZymes, overall, more than half of the CAZyme-containing BACs presented multiple CAZymes in tandem with important regulators, transporters, and other genes with a variety of functions (Figs. 3 and 4). Careful inspection of such clusters must be conducted to improve the understanding of the coregulatory patterns and synergistic actions among clustered genes, clarifying the lignocellulolytic activities of *Trichoderma* ssp.

### Expression measurement and DEG identification

A prevalidated RNA-Seq (Almeida et al., 2021) dataset was used as a source to identify regions/genes potentially activated during lignocellulose degradation. These previous studies characterized the global transcriptome through transcriptomic profiling, revealing different strategies for carbon utilization (Almeida et al., 2021). Transcriptomic, genomic, and phenotypic variation is common even at the strain level, and such differences must be carefully studied to understand their role in the biodegradation process (Almeida et al., 2021; Hewedy et al., 2020; Pachauri et al., 2023; Rosolen et al., 2023). In TA0020 and TH0179, genes exhibited significant differences in expression, but few genes showed significant values in the differential expression analysis. These genes are likely regulated in response to the studied conditions; thus, these studies show the genomic context of these genes (Figs. 3 and 4).

The complex transcriptomic profile of TA0020, observed in the previous *de novo* transcriptome analysis, was captured in this reference-based transcriptome assembly and differential expression analysis. Their lower lignocellulolytic potential contrasts with their complex transcriptome profile, likely because *T. atroviride* may not be as specialized in lignocellulose utilization as the saprophytic workhorse *T. reesei* QM6a or the *T. harzianum* IOC 3844 and *T. harzianum* CBMAI-0179 strains, which show greater efficiency in crystalline cellulose degradation (Almeida et al., 2021; Horta et al., 2018; Li and Wang, 2021).

## Conclusions

The BAC methodology allowed us to recover CAZyme-containing genomic regions and study their linear distribution, genomic context, and relationship to degradative conditions. It was possible to identify target lignocellulolytic CAZyme clusters and to address the differential lignocellulolytic potential in terms of gene expression previously observed among species. Functional inference combined with transcript expression quantification and comparative analysis provided highly valuable information regarding the genome arrangement of CAZyme genes, making it possible to identify strain-specific characteristics. We recovered multiple CAZyme-containing regions rich in TF- and TR-flanking genes, revealing the control mechanisms for CAZyme expression. Expression analysis helped to identify regions that were positively regulated under biomass degradation conditions and likely involved in the lignocellulolytic activities of both strains.

These results contribute to the genomic perspective of targeted CAZyme regions in nonmodel *Trichoderma* species to identify potential highly degradative regions, which is useful for improving enzymatic degradation technologies. Here, we provided an in-depth analysis of the target lignocellulolytic genomic regions of novel *Trichoderma* strains.

## Supporting information

Supplementary_material_overview

## Acknowledgements

A special thanks to CPQBA-CBMAI for providing the fungi samples and to CBMEG for providing high-quality infrastructure to accomplish the studies.

## Declaration of generative AI and AI-assisted technologies in the writing process

No AI-based technology was used in the writing process.

## Data availability

All of the data generated or analyzed in this study are included in this published article (and its supplementary information files). The assembled genomic regions were deposited in the NCBI database via BankIt and can be accessed under BioProject numbers PRJNA1028979, TH0179 SAMN37846728 and TA0020 SAMN37846658.

## Author contributions

PHC: writing—original draft, methodology, formal analysis, and visualization. RRR: conceptualization, methodology, and writing—review & editing. MACH: conceptualization, methodology, and writing—review & editing. JSM: methodology and resources. CCS: methodology and resources. DAS: methodology and resources. APS: conceptualization, supervision, review & editing, and funding acquisition.

RRR: MACH: PHCA: methodology and resources. CCS: methodology and formal analysis. DAS: GHG: writing—review & editing.

## Abbreviations

BCA: Biocontrol agents
CAZyme: Carbohydrate-active enzymes
TF: Transcription factor
TR: Transporters
SMBGC: Secondary metabolism biosynthesis-related gene cluster
BAC: Bacterial artificial chromosome
PDA: Potato dextrose agar
DEG: Differential expression of genes

